# DiscoEPG: A Python package for characterization of insect electrical penetration graph (EPG) signals

**DOI:** 10.1101/2024.12.05.627099

**Authors:** Quang Dung Dinh, Daniel Kunk, Truong Son Hy, Nalam Vamsi, Phuong D. Dao

**Affiliations:** Institut Galilée, Universite Sorbonne Paris Nord, Villetaneuse 93430, Paris, France; Department of Agricultural Biology, Colorado State University, Fort Collins, CO 80523, United States; Department of Cell and Molecular Biology, Colorado State University, Fort Collins, CO 80523, United States; Graduate Degree Program in Ecology, Colorado State University, Fort Collins, CO 80523, United States; Department of Computer Science, University of Alabama at Birmingham, Birmingham, AL 35294, United States

**Keywords:** Electrical penetration graph, Pierce-sucking insect, Automatic annotation, Machine learning, Open-source package

## Abstract

The Electrical Penetration Graph (EPG) technique is a well-known specialized tool that entomologists use to monitor and analyze the feeding behavior of piercing-sucking insects, such as aphids, whiteflies, and leafhoppers, on plants. Traditionally, the annotation is conducted by a well-trained technician who uses expert knowledge to compare the targeted waveforms with standard waveforms of aphid feeding behavior, which takes approximately 30 minutes to annotate an 8-hour recording depending on the complexity of the insect behaviors. Machine learning (ML) models, which shown great potential in monitoring insects behaviors, have recently been used to speed up this process. However, most publicly available tools that provide automatic annotation suffer from low prediction accuracy due to only using simple distinction rules to classify waveforms. For this reason, we develop *DiscoEPG*, an open-source Python package which performs accurate automatic EPG signal annotation. Various ML algorithms were experimented rigorously, which reports greater prediction power and improved accuracy in comparison to previous studies. In addition, we equipped our package with novel tools for generating journal-level plots to facilitate visual inspection, while including the computation of various EPG parameters and necessary statistical analysis which are popular in the research of aphids. With *DiscoEPG*, we aim to facilitate the rapid characterization of analysis of piercing-sucking insects feeding behavior through EPG signal, making this technique more viable to researchers who share the same interest. Our package is publicly available at: https://github.com/HySonLab/ML4Insects.

## 1 Introduction

First proposed in 1964 by McLean and Kinsey [1] and later developed by Tjallingii and Hogen Esch [2], the electrical penetration graph (EPG) technique has been used to monitor the feeding behavior of hemipteran insects that feed on plant phloem sap using their needle-like mouthparts. Since then, this technique has become essential for understanding the interactions between various species of insects and their host plants, and more than 50 species have been successfully studied. The technique works by creating a closed electrical circuit that connects individual insects to their host plant, followed by recording the amplified voltage changes that are generated as the insect performs feeding activities by puncturing into the plant tissues. According to Tjallingii and Hogen Esch [2], distinctive waveform patterns observed from EPG recordings reflect different actions performed by the insect stylet, which correspond to different feeding behaviors. The EPG technique has been used to investigate many facets of plant-insect interactions, including the identification and characterization of host plant resistance [3], the transmission mechanisms of plant pathogens by their insect vectors [4], and the mode of action of pesticides [5]. Computer programs such as *Stylet+* [6] were developed to transform EPG data into digital formats with user-friendly and mouse-driven interfaces, while providing additional utilities for visualizing EPG signals at scale and manually labeling EPG waveforms. Once waveforms are identified and annotated, insights into plant-insect interactions are yielded by analyzing the insect feeding behaviors using a range of computed parameters and statistical tests. Despite its power, the feasibility of the EPG technique is greatly limited by the time and complexity of the manual annotation and parameter estimation processes. For example, an experienced observer may need about 30 minutes to thoroughly annotate a recording, depending on the length and the complexity of the observed waveforms. Additionally, manual waveform annotation can be prone to human error, and variability in waveform calls can arise between annotators. Therefore, it is critical to develop an accurate and automated computer program to overcome these obstacles and thus make the EPG technique more robust and accessible to researchers. In fact, various studies have proposed tools for this purpose, as mentioned in Table 1.

**Table 1:**
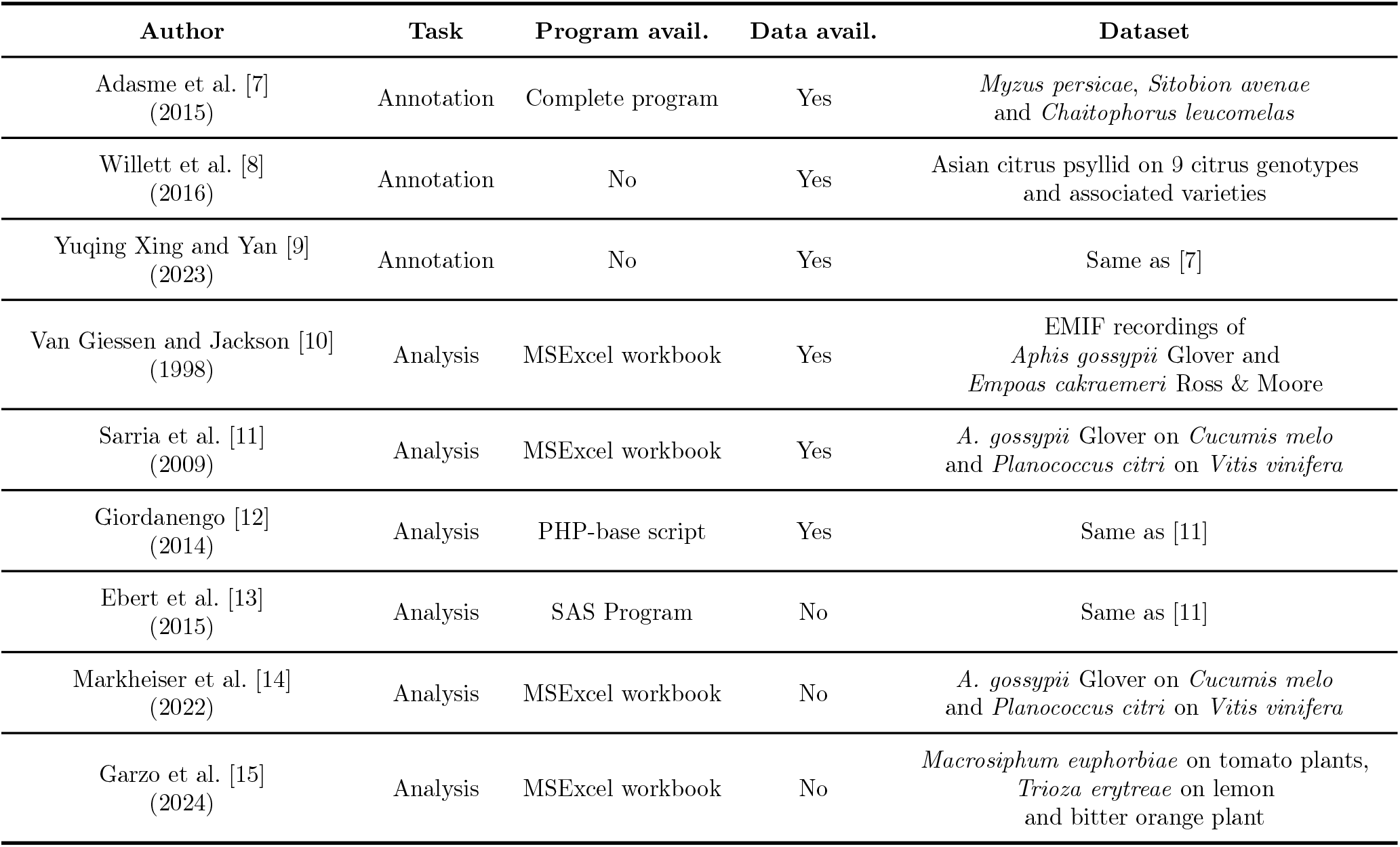
Some recent studies on automating EPG analysis tasks, including annotation (labeling an aphid’s behavior) and analysis (calculating EPG parameters and performing statistical tests).

| Author | Task | Program avail. | Data avail. | Dataset |
| --- | --- | --- | --- | --- |
| Adasme et al. [7]<br>(2015) | Annotation | Complete program | Yes | *Myzus persicae*, *Sitobion avenae*<br>and *Chaitophorus leucomelas* |
| Willett et al. [8]<br>(2016) | Annotation | No | Yes | Asian citrus psyllid on 9 citrus genotypes<br>and associated varieties |
| Yuqing Xing and Yan [9]<br>(2023) | Annotation | No | Yes | Same as [7] |
| Van Giessen and Jackson [10]<br>(1998) | Analysis | MSExcel workbook | Yes | EMIF recordings of<br>*Aphis gossypii* Glover and<br>*Empoas cakraemeri* Ross & Moore |
| Sarria et al. [11]<br>(2009) | Analysis | MSExcel workbook | Yes | *A. gossypii* Glover on *Cucumis melo*<br>and *Planococcus citri* on *Vitis vinifera* |
| Giordanengo [12]<br>(2014) | Analysis | PHP-base script | Yes | Same as [11] |
| Ebert et al. [13]<br>(2015) | Analysis | SAS Program | No | Same as [11] |
| Markheiser et al. [14]<br>(2022) | Analysis | MSExcel workbook | No | *A. gossypii* Glover on *Cucumis melo*<br>and *Planococcus citri* on *Vitis vinifera* |
| Garzo et al. [15]<br>(2024) | Analysis | MSExcel workbook | No | *Macrosiphum euphorbiae* on tomato plants,<br>*Trioza erytreae* on lemon<br>and bitter orange plant |

From Table 1, a greater proportion of works [10, 11, 12, 13, 14, 15] is devoted for computer programs that quickly calculate the behavioral parameters and statistics of EPG data, which were built on different environment such as PHP, SAS or MSExcel. Although these programs support analyses of various parameters, they suffer from limitations such as having calculation errors or not being able to process data in batches. The latest publication [15] addresses these limitations with their MSExcel workbook, which support a set of parameters with explicit definitions suitable for studying the behaviors of aphids and psyllids. The other groups offer solutions for speeding up the EPG waveform annotation process. Multiple approaches have been used, including using direct numerical constraints, or leveraging machine learning algorithms. For example, Adasme et al. [7] presented a search engine that used a set of numerical constraints computed from the amplitudes of the EPG signal to distinguish the feeding stages of aphids, namely NP, C, E1, E2, F, G and pd. Due to the overly restrictive constraints, the method suffers from low classification performance. Meanwhile, Willett et al. [8] used machine learning algorithms to discover the relationship between the virus transmission from the Asian citrus psyllid (*Diaporina citri*) and different genotypes of citrus plants. They achieved a high overall classification accuracy (97.4%) with Random Forest, while also performing hierarchical clustering to explore the waveform similarity between feeding stages. Nevertheless, experiments in these works were performed on relatively small scales, and more importantly, no well-documented programs were published, making it challenging for users to verify the reported result and re-implement such approaches in new experiments.

To fill the gaps and resolve the outstanding issues identified in the previous works, we have developed an open-source Python package, DiscoEPG, designed to aid in EPG analysis. This innovative toolkit is designed to efficiently import data obtained from *Stylet+*, while offering a comprehensive set of convenient functions for characterizing aphid feeding behavior. The major contribution of our package is presented by the high-quality and automatic annotation of EPG signal using ML algorithms. 6 well-established algorithms from both the traditional ML group (e.g., Extreme Gradient Boosting, Random Forest) and the thriving deep learning group (e.g., Convolutional Neural Networks) on across different experimental settings. The annotation results can be easily refined and completed with *Stylet+* to eliminate any imperfections in annotation from the ML model. These experiments also include an ablation study on feature extraction techniques, where we learn to determine the inputs that generate the best annotation result [16]. In comparison to previous studies, our model evaluations were carried out rigorously at a much larger scale and achieved comparative metrics. To facilitate the investigation insect feeding behavior, DiscoEPG is equipped with visualization modules which generate not only publication-standard plots, but also interactive plots where zooming and navigating waveforms is possible within mouse clicks. Lastly, the package offers automatic calculation of EPG parameters and statistical test (Students t-test, Wilcoxon test, ANOVA and Kruskal Wallis) that assist with in-depth analysis of aphid feeding behavior. Although initially developed based on aphid data, our package can be easily extended to work with EPG data from other hemipteran insects.

## 2 Development of DiscoEPG

### 2.1 Electrical Penetration Graph (EPG) data

For aphids, their feeding activities create specific patterns which are generally termed *waveforms* [17, 18, 19, 20, 21, 22]. These activities are divided into seven categories including non-probing phase (NP), pathway phase (C), potential drop (pd), xylem ingestion (G), penetration difficulties (F), salivation (E1) and ingestion (E2) [23], based on their amplitude and frequency. By analyzing various parameters related to these waveforms (e.g. occurrence frequency, total length, average length), researchers gain insights into various aspects of aphid feeding behavior and thus enable evaluation of the impact of aphid feeding on plant health.

For testing DiscoEPG, our experiments were conducted on 5 datasets recorded a large and diverse set of datasets obtained from 4 species of aphids and 6 species of host plants (8 varieties), spanning over a total of 332 8-hour recordings: *Rhopalosiphum padi* (bird cherry-oat aphid; BCOA1, BCOA2), *Myzus persicae* (green peach aphid; GPA), *Phorodon cannabis* (cannabis aphid; CA), and *Aphis glycines* (Soybean aphid; SA). Recent works providing automated tools for EPG analysis were conducted on much smaller datasets, making our work one of the most comprehensive studies so far. For a detailed description of each dataset, please refer to [16].

### 2.2 Package development

To the best of our knowledge, DiscoEPG is the first multi-functional open-source package for facilitating aphid feeding behavior analysis. Our first and main goal for developing DiscoEPG was to build a well-documented package which allows users to easily reproduce our experiments on the application of multiple machine learning algorithms for automatically conducting EPG waveform annotation. DiscoEPG was developed solely in the Python programming language, where the ML algorithms used in the package are implemented based on well-known Python libraries such as *scikit-learn* and *PyTorch*. The plotting utilities were made with the *plotly* library, which is a well-known Python library for making interactive plots. In order to effectively visualize a large amount of data points (since each EPG recording may contain millions), we utilize another library called *plotly-resampler* [24], which is a toolkit that supports memory-efficient data visualization. The syntax of DiscoEPG’s functions, which will be discussed further in Section 3, are written in simple language and easy to understand. All of our experiments were implemented on a single laptop GIGABYTE G7, with an Intel Core i5-10500H CPU (12 cores), 6GB of memory and an NVIDIA GeForce RTX 3060 Laptop GPU. Our package can be run on common personal computers, although a computer with a GPU of equivalent or higher specifications is recommended to ensure smooth implementation of deep learning architectures.

## 3 Main components of DiscoEPG

### 3.1 *EPGDataset*: Reading, displaying and analyzing EPG data

#### Reading EPG data

In DiscoEPG, we constructed an object class that is dedicated to reading, displaying, and analyzing EPG data, namely *EPGDataset*. Table 2 describes some of the main functions of *EPGDataset*. Generally, each recording comprises of several hour-long recording files (in *.D0x extension) in a binary encoding format along with a text file that provides corresponding ground-truth annotations (in *.ANA extension). In our experiments, each recording was sampled at 100Hz for 8 hours (8 *.D0x files total per recording), resulting in a single signal channel with 2,880,000 time steps. *EPGDataset* is able to quickly read a directory with hundreds of hours of EPG recordings in seconds (we read 332 *.D0x files in under 10 seconds). 11 guidelines [15] will be automatically checked and results will be exported into an *.xlsx file in the working directory.

**Table 2:**
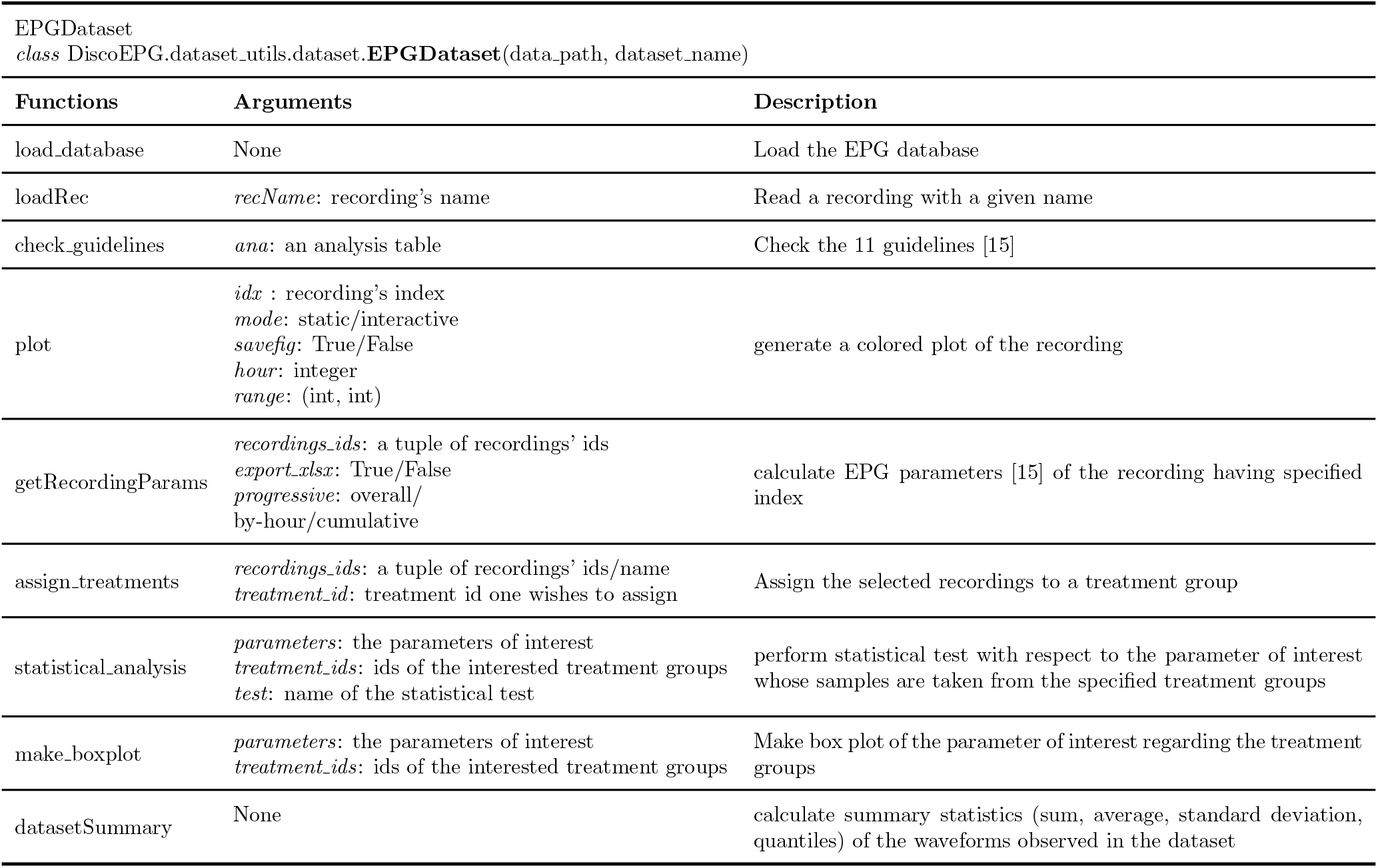
Some main functions of *EPGDataset*.

#### EPG parameters calculation and statistical analysis

The function *EPGDataset*.*getRecordingParams* provides a rapid solution for the task of calculating feeding behavior parameters from EPG data. Thanks to the ability to read multiple recordings at the same time, DiscoEPG is capable of rapidly computing all parameters across the entire experimental duration for multiple recordings at once. To gain additional insights into how behaviors change over time, users can provide additional instruction to this function through the *progressive* arguments by setting it to either “overall”, “by hour” or “cumulative”. The “overall” options shows the parameters over an entire recording duration. Meanwhile, the “by hour” option calculates a subset of variables specific to each hour, and the “cumulative” option tracks these variables over time, from the start of the recording through the first, second, third hour, and up to the end. DiscoEPG is able to output these results into a Microsoft Excel *.xlsx file for further inspection. In terms of units, we use seconds for all parameters, regardless of the waveform duration. The program outputs a log file showing the errors detected during calculation (such as the absence of a waveform, incorrect format, etc).

#### Data visualization

DiscoEPG is equipped with various functions for making aesthetic plots of the EPG signal in both static and interactive modes. With interactive plots, users can easily engage with the figures through zooming, sliding, and more; enabling direct monitoring of EPG signals within their Jupyter Notebook environment. This serves as a quick visual inspection tool allowing users to get an overview of feeding behavior patterns and their locations, which is highly beneficial for post-prediction processing and error correction. Rather than plotting the whole recording at once, users may also choose to restrict the plot to specific intervals with the two arguments *hour* and *range*. In addition, it is possible to visualize the automatic segmentation results side-by-side with the ground-truth annotation in order to visually assess the performance of each machine learning algorithm.

### 3.2 *EPGSegment* and *EPGSegmentML*: Automatic annotation of EPG waveforms

#### Method

In DiscoEPG, we introduce for the first time a novel feature for predicting waveform locations using state-of-the-art ML algorithms. Six well-established ML models were studied for the classification task, which were divided into the traditional group (supported by the *EPGSegment* class), and the deep learning group (supported by the *EPGSegmentML* class). The explicit descriptions of the functions of *EPGSegment* and *EPGSegmentML* are given in Table 3 and Table 4, respectively. For the former group, Random Forest (RF), Extreme gradient boosting (XGB) and Logistic Regression (LR) were implemented, in which the first two demonstrated excellent waveform recognition ability. Meanwhile, the latter group contains three models from the family of convolutional neural networks (CNNs), namely 1DCNN, ResNet (operating on one dimensional input) and 2DCNN (operating on two dimensional input). We developed a three-stage pipeline for automating the annotation process, which is composed of the initial segmentation stage, the waveform sample classification stage, and the label aggregation stage, as illustrated in Fig 1. In the first stage, an input signal is preprocessed by dividing into consecutive segments whose are of uniform length of *d* time steps (*d* = 1024 by default, corresponding to 10.24s), then applying signal transformation techniques (e.g. Fourier transform) to extract features in the frequency domain. Then, ML models are used to classify such segments using the transformed features, which assign to a segment the label corresponding to the highest probability. The final stage is dedicated to aggregating these predictions in order to obtain a complete annotation for the input recording, from which one may determine the predicted locations of each waveform. The result of the automatic annotation process is an analysis text file (*.ANA) saving the positions of each waveform.

**Table 3:** Some main functions of *EPGSegment*.

| EPGSegment<br>*class* DiscoEPG.models.Segmentation.**EPGSegment**(config, inference = False) |  |  |
| --- | --- | --- |
| Functions | Arguments | Description |
| train | *early_stop*: True/False<br>*patience*: integer<br>*min_delta*: float<br>*verbose*: True/False | train the model with the specified dataset. Arguments allow the training process to stop in case the validation accuracy does not surpass a minimum threshold. |
| evaluate | *task*: string: val/test | Evaluate the Task 1 (classification) performance of the trained model |
| load_checkpoint | *path*: string | load a trained model |
| save_checkpoint | *name*: string | load/save the trained model |
| plot_train_result | *savefig*: True/False | Display the training curves and evaluation metrics of Task 1 |
| segment | *recording_name*: string | Predict a complete segmentation for the recording |
| save_analysis | *name*: string | Save the predicted annotation |
| plot_segmentation | None | Plot the predicted and ground-truth segmentation side-by-side |

**Table 4:** Some main functions of *EPGSegmentML*.

| EPGSegmentML<br>*class* DiscoEPG.models.Segmentation.**EPGSegmentML**(config, inference = False) |  |  |
| --- | --- | --- |
| Functions | Arguments | Description |
| get_traindata | *test_size*: float | preprocess and split the dataset into train and test set, then perform feature extraction |
| fit | *X_train*: np.ndarray<br>*y_train*: np.ndarray | train the ML model |
| predict | *X_test*: np.ndarray<br>*y_test*: np.ndarray | perform classification on the test set for Task 1 evaluation |
| plot_train_result | *savefig*: True/False | Display the evaluation metrics of Task 1 |
| segment | *recording_name*: string | Predict a complete segmentation for the recording |
| save_analysis | *name*: string | Save the predicted annotation |
| plot_segmentation | None | Plot the predicted and ground-truth segmentation side-by-side |

**Figure 1:**
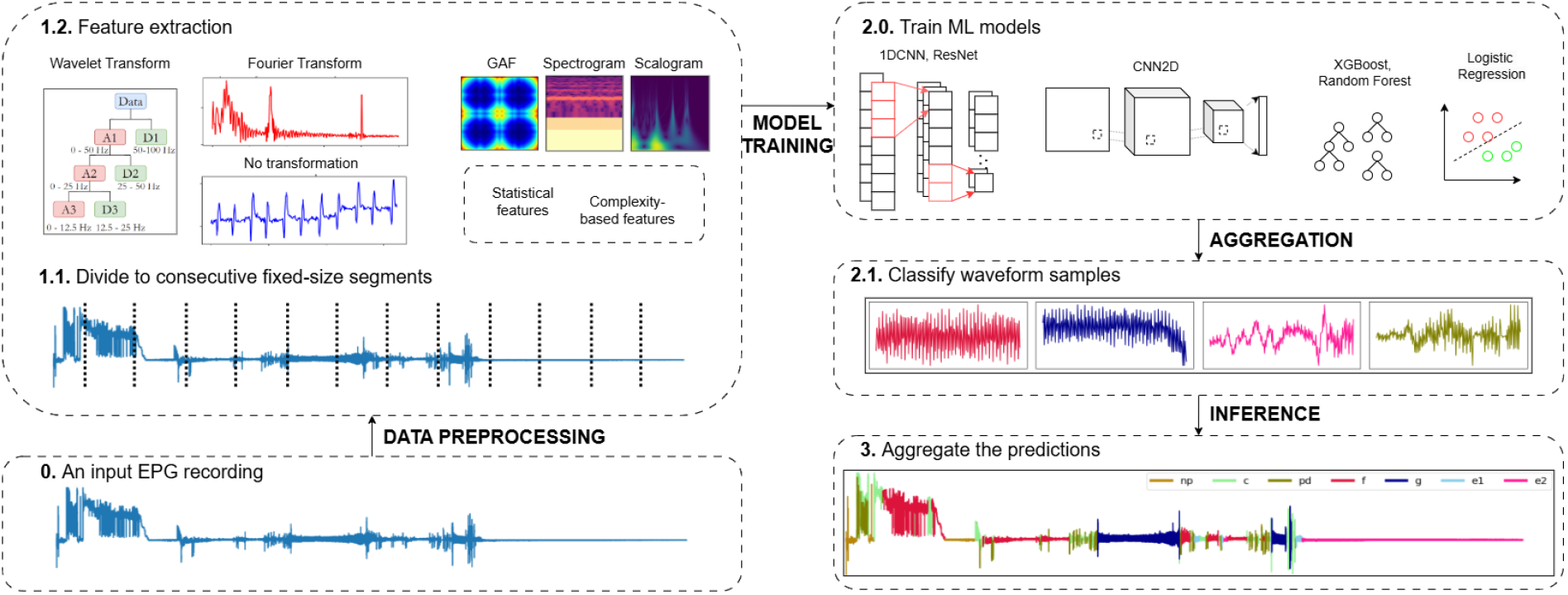
Our proposed pipeline for applying ML method to automatically annotate EPG data. In the initial stage, the recording are segmented into consecutive segments of uniform length and then preprocessed using signal transformation technique. Next, ML models are trained with the transformed data to classify the obtained waveform samples. Finally, the predictions are aggregated, yielding a complete annotation, from which the user can determine the predicted location of each waveform.

#### Feature extraction

Previous studies on the time-series adaptation of RF [25, 26] suggested classifying a time-series using a range of features calculated from its component segments. In this process, simple statistical features such as mean, standard deviation and slope were found to be computationally efficient, outperforming strong competitors such as 1-NN classifiers with dynamic time-warping distance. This idea was adopted for our study, in which we consider 13 statistical and complexity features, namely the mean, root mean square, standard deviation, variance, skewness, quantiles at levels 0.05, 0.25, 0.5, 0.75, 0.95, zero crossing rates, Shannon entropy, and permutation entropy of the consecutive segments for training traditional machine learning algorithms. Regarding the deep learning algorithms, feature extraction is not required since the characteristic features are learned autonomously through a huge array of parameters. Therefore, we train the CNNs directly on the frequency domain information. For each model, we also tested the effect of three data input types including no transformation, Fourier transform, and wavelet transform. The results show that no transformation and Wavelet transformation inputs yield the best performance for 1DCNN and XGB, respectively.

#### Evaluation

To thoroughly evaluate the performance of our proposed pipeline, we conducted two-fold assessment regarding the classification performance of ML models and the localization performance of the sliding window strategy (Fig 1). For the former task, per-class and global metrics were taken into account, including the per-class accuracy score, the overall accuracy (OA) score and the average F1 (avg-F1). Here, the average F1 score is the average of the F1 scores of all fixed-length waveform instances. These scores essentially tell us how well the ML models are able to classify insects’ feeding behaviors based on the observed pattern. The formulas for Accuracy and F1 metrics are given by:

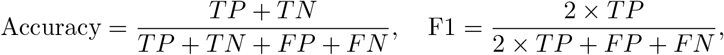

where the acronyms TP, TN, FP, and FN refer to True Positive, True Negative, False Positive, and False Negative, respectively. For the latter step, we consider the overlap rate (OR), which is the ratio at which the predicted waveform positions coincide with the ground-truth waveform positions. This metric aligns well with our ultimate goal, which is to perform the detection of insects’ behavior in an EPG recording.

## 4 Results

### 4.1 Automatic segmentation results

In this section, we select and report the results of XGB and 1DCNN models, which are the representative candidates from the traditional ML group and the deep learning group, respectively. The selection was based on both performance and computational complexity, as each demonstrated competitive metrics along with fast training and inference speeds. Validation was conducted on a large dataset to evaluate the models’ generalization across different scenarios. XGB and 1DCNN describe strong performance in automatic EPG signal annotation and characterization, achieving an OA of over 90% in the classification task. Fig 2 provides a detailed description of the evaluated metrics. The per-class classification accuracy scores ranged from 84% to 98%, demonstrating the potential of using ML algorithms for characterizing aphids’ feeding stages. Among the different feeding behaviors, NP, E2, F and G demonstrate relatively higher annotation accuracy thanks to their distinctive patterns. In contrast, it was challenging to distinguish between C and pd since C is a complex waveform which is usually composed of a mixture of patterns while pd is a waveform contained within C. By analyzing the confusion matrices, we found that C and pd are usually misclassified with one another. Finally, the OR for XGB and 1DCNN, which display the waveform localization performance, were 80.6% and 75.7%, respectively.

**Figure 2:**
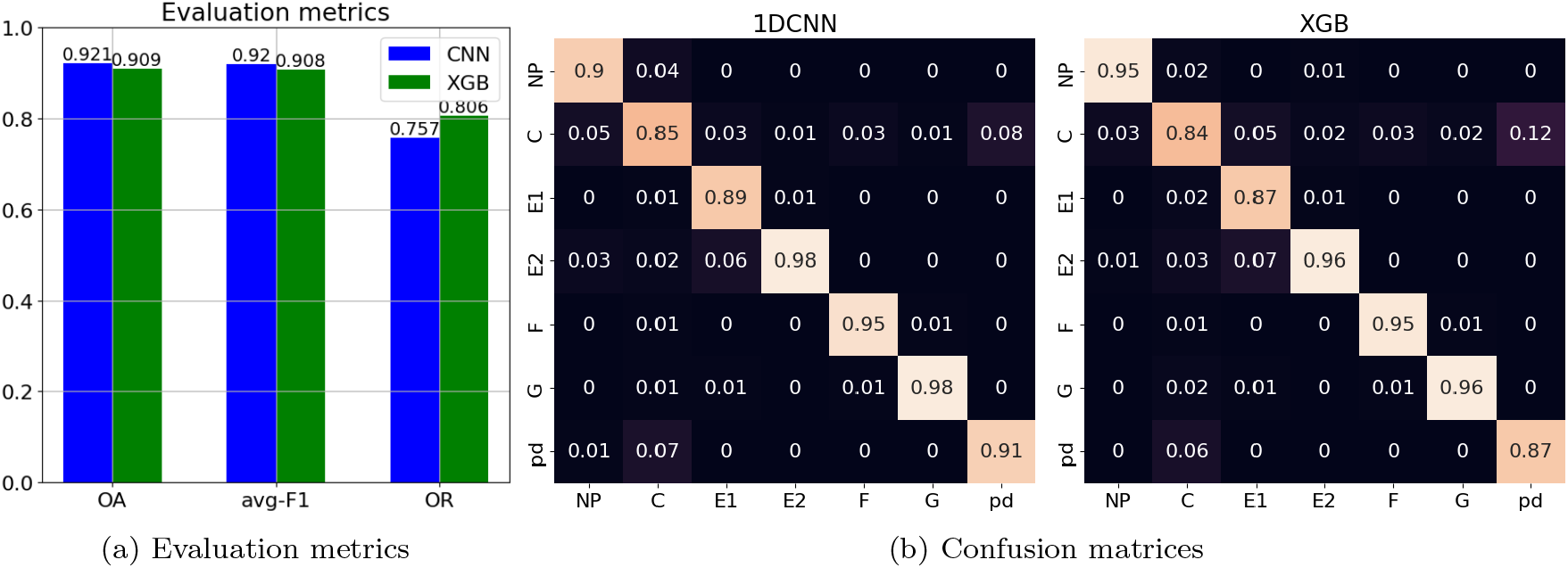
Evaluation metrics of 1DCNN (left) and XGB (right) and confusion matrices of two representative machine learning models, trained on a combined dataset.

In Fig 3, we present an example of the prediction made by DiscoEPG (lower) in comparison to the ground-truth annotation (upper) in which the reported OR was 90.18%. Most of the time, the long and dominant waveforms such as G and E2 are easily detected, while it is more challenging for the model to detect C, a complex waveform as mentioned. The situation is even more troublesome with short waveforms such as pd. The segment created by the sliding window technique often overlook the existence of these waveform as they have fixed end points, such as from time steps 0 to 1023, 1024 to 2047, 2048 to 3071 and so on. Thus, as observed in Fig 3b which is a zoomed-in version of Fig 3a from 7000s to 8000s, we found that misclassification of pd to C are visible. Besides, the misalignment between endpoints of waveform can also be attributed to the same reason. Therefore, it is recommended that users perform a post-prediction assessment to avoid these inaccuracies.

**Figure 3:**
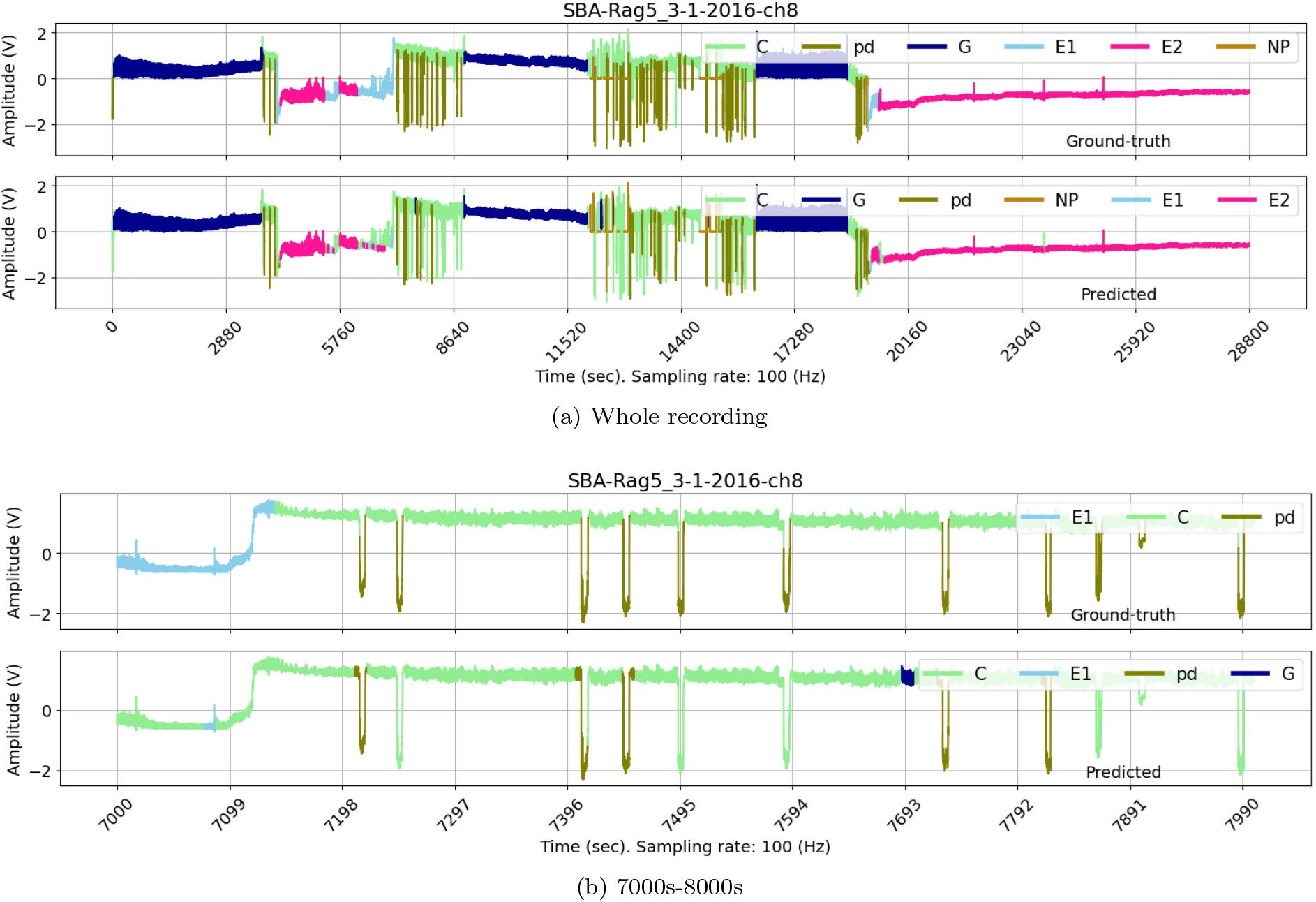
Ground-truth vs. predicted annotation of a recording taken from the SBA dataset. Sub-figure a) and b) displays the comparison over the whole recording and the period 7000s-8000s, respectively.

### 4.2 Example of usage

In our code, provided at https://github.com/HySonLab/ML4Insects, we provide example Jupyter notebooks that explain step-by-step the procedure for training ML model and making prediction as well as the procedure to perform behavioral parameter calculations and statistical test on the parameters.

For data analysis, we refer the user to the *EPGDataset* class, which supports loading EPG recordings, visualizing EPG data, and performing EPG analysis. *EPGDataset* takes two input: the path to the data folder, and the name of the dataset. The EPG recordings will be loaded as soon as the class *EPGDataset* is called. Visualization can be done with the functions *plot*, which generates a figure of the specified recording in either static or interactive mode. For the interactive mode, an additional argument *smoothen* can be used, stating if *plotly-resampler* can be used to smoothen the data for a memory-efficient plot, i.e. one which is generated faster and consumes less memory. Before performing analysis, it is recommended that users divide recordings into treatment groups, which can be done using the function *assign_treatments*. Next, calling the function *getRecordingParams* will automatically compute the behavioral parameters as mentioned in Section 3.2 for in-depth EPG analysis. Users can choose to work with any of the loaded recordings by providing a list of integers for the *list_of_indices* argument, and to export the calculated results to an *.xlsx file with the *export_xlsx* argument. Moreover, calculated results can be displayed in either row or column format by adjusting the argument *view*. After assigning treatment groups and calculate the EPG parameters, the users may conduct statistical tests with the function *statistical_analysis* by simply inputing the parameters, the treatment groups and the test of interest. Algorithm 1 provides a snippet of implementing the main functions in a Python environment.

#### Algorithm 1: Performing parameters calculation and data visualization with DiscoEPG.

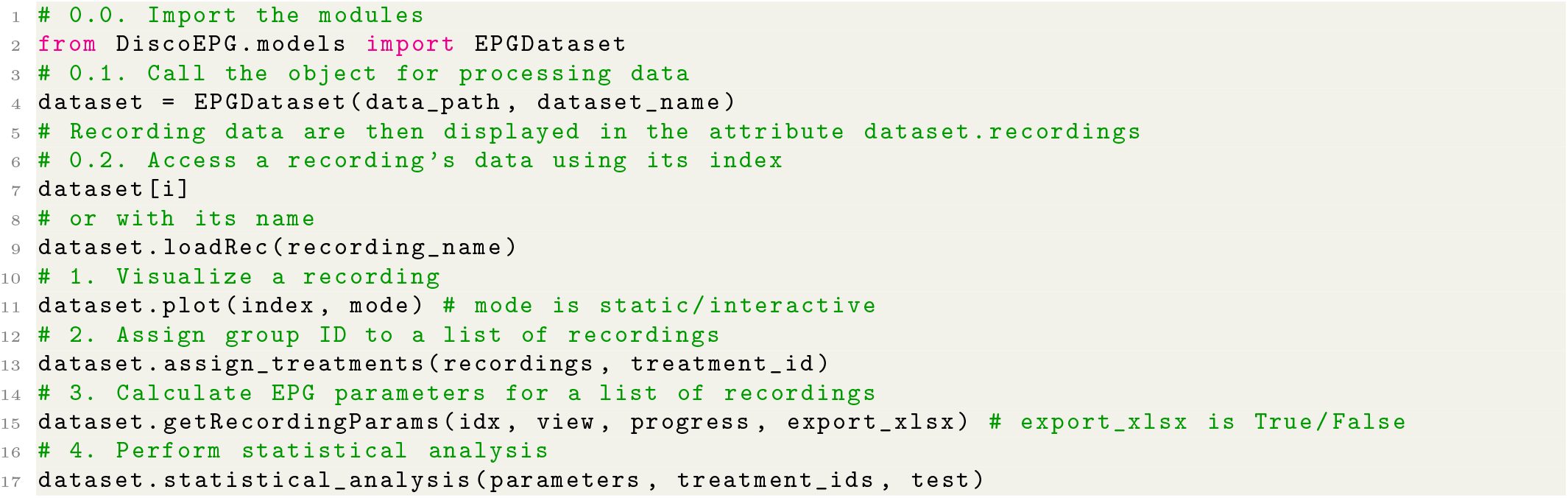

To work with ML models, two classes *EPGsegment* and *EPGsegment ML* support training and inference with deep learning and traditional ML models, respectively. We present an example of using *EPGsegment* for simplicity as the procedures are quite similar. Algorithm 2 provides an example of performing this pipeline with DiscoEPG. First, a configuration file in *\**.*json* format is often used to save the configurations to the EPG database and the settings of the ML models. It must be processed into a Python dictionary by the *process_config* utility function and used as input for *EPGsegment*. Upon calling *EPGsegment*, the trainer automatically retrieves the EPG database, performing data preprocessing and gets ready for the training/inference process. Training the ML models is simple by calling the function *train*. Then, evaluation of the classification of waveform samples can be done immediately. After that, the model can be easily saved with the function *save_checkpoint* for later use. Automatic segmentation function is provided by the function *segment*, which takes the recording’s name, or the recording’s ID as input argument. Finally, users can visually investigate the quality of the predicted segmentation with the *plot segmentation* function. Note that, this function also adopts the arguments from the basic *plot* function from *EPG*.*Dataset*, so the users can freely change the plots’ sizes, ratios or the duration to plot. In the case that the recording has a ground-truth annotation, the overlap rate (OR) metric for the predicted annotation and the ground-truth annotation is displayed along with the function *segment*.

#### Algorithm 2: Simple syntax for training a ML model and making inference with DiscoEPG in Python style.

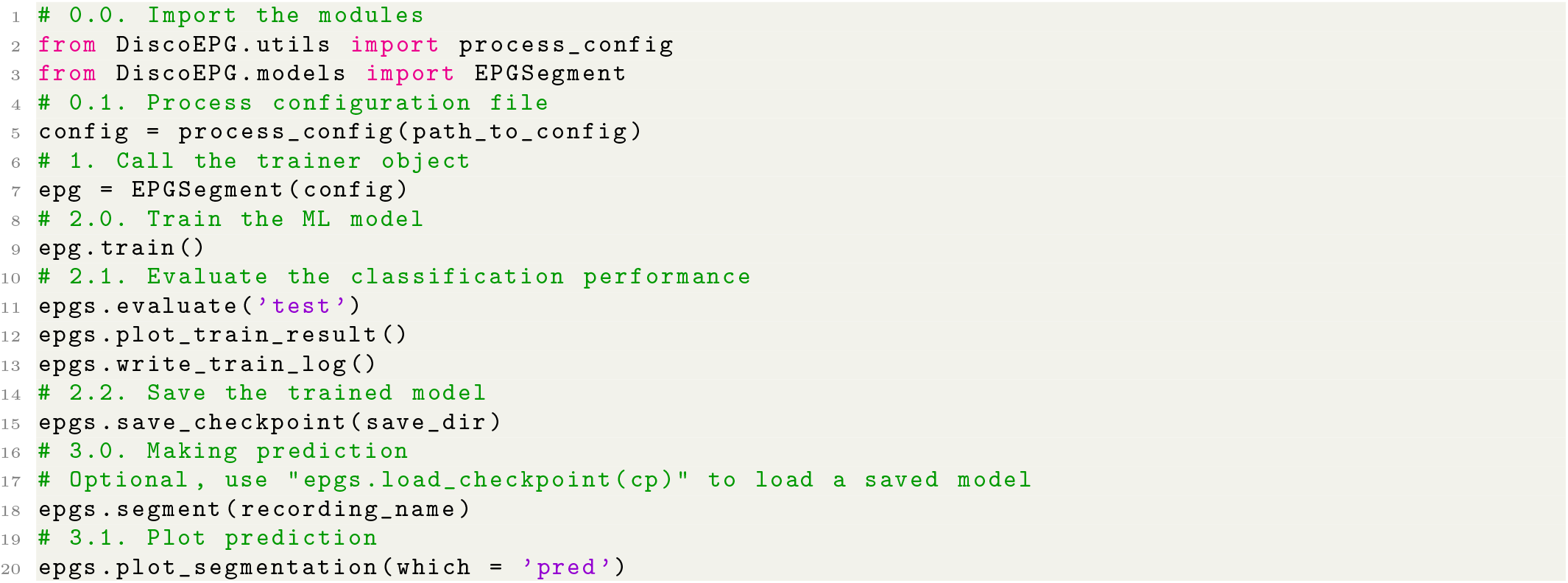

## 5 Discussion

In this paper, we present an open-source Python package, DiscoEPG, as an advanced and effective tool to assist with the analysis of aphid feeding behavior through the EPG technique. DiscoEPG provides three main functions: EPG signal reading and visualization, automatic waveform annotation, and EPG behavioral parameter calculation. Notably, we introduce for the first time the use of modern machine learning algorithms for characterizing EPG data, which yield comparative results with previous studies, but with much more flexibility in model parameter settings. With these functions, DiscoEPG seeks to address the bottlenecks related to the time and computational complexity of EPG data analysis, cutting annotation time from hours to just minutes for projects with large datasets. Despite various advantages, our package demonstrates several limitations that can be further enhanced in future studies. Regarding the automatic waveform annotation process, the sliding-window technique creates fixed waveform segments as inputs of ML models while actual waveform segments of different feeding behaviors can vary. Therefore, a more flexible segmentation algorithm is recommended to generate more representative labeled data for the ML models. One possible idea is to construct a neural networks that automatically learns the waveforms position, as detailed in UTime, an adoption of a fully convolutional neural network for segmentation named UNet. Empirically, we found that neural network classification models shows robust performance in recognizing EPG waveforms, while also being advantageous for eliminating the need for the feature extraction process. Hence, studying other deep neural network architectures promises to further improve our results. Software such as *Stylet+* can provide better graphical representations that allow interactive scaling and zooming features, which can show more fine details than our package. Overall, DiscoEPG provides researchers with a reliable and user-friendly tool for automatically annotating and characterizing EPG data to help understand the feeding behavior of piercing-sucking insects.

## References

[1] DL McLean and MG Kinsey. A technique for electronically recording aphid feeding and salivation. Nature, 202:1358–1359, 1964. doi: 10.1111/j.1365-3032.1993.tb00604.x.

[2] W.F. Tjallingii and Th. Hogen Esch. Fine structure of aphid stylet routes in plant tissues in correlation with epg signals. Physiological Entomology, 18:317–328, 1993. ISSN 0307-6962. doi: 10.1111/j.1365-3032.1993.tb00604.x.

[3] Carmen Escudero-Martinez, Daniel J Leybourne, and Jorunn IB Bos. Plant resistance in different cell layers affects aphid probing and feeding behaviour during non-host and poor-host interactions. Bulletin of Entomological Research, 111(1):31–38, 2021.

[4] Wenbin Lei, Pei Li, Yongqiang Han, Shaolong Gong, Lang Yang, and Maolin Hou. Epg recordings reveal differential feeding behaviors in sogatella furcifera in response to plant virus infection and transmission success. Scientific reports, 6(1):30240, 2016.

[5] Yueping He, Li Chen, Jianming Chen, Juefeng Zhang, Liezhong Chen, Jinliang Shen, and Yu Cheng Zhu. Electrical penetration graph evidence that pymetrozine toxicity to the rice brown planthopper is by inhibition of phloem feeding. Pest Management Science, 67(4):483–491, 2011.

[6] Vicenta Salvador Recatala and Fred Tjallingii. A new application of the electrical penetration graph (epg) for acquiring and measuring electrical signals in phloem sieve elements. Journal of visualized experiments : JoVE, 2015, 07 2015. doi: 10.3791/52826.

[7] Francisco Adasme, Camila Muñoz, Josselyn Salinas-Cornejo, and Claudio Ramírez. A2epg: A new software for the analysis of electrical penetration graphs to study plant probing behaviour of hemipteran insects. Computers and Electronics in Agriculture, 113:128–135, 02 2015. doi: 10.1016/j.compag.2015.02.005.

[8] Denis Willett, Justin George, Nora Willett, Lukasz Stelinski, and Stephen Lapointe. Machine learning for characterization of insect vector feeding. PLOS Computational Biology, 12:e1005158, 11 2016. doi: 10.1371/journal.pcbi.1005158.

[9] Lili Wu Yuqing Xing, Baofang Li and Fengming Yan. Waveforms eavesdropping prevention framework: The case of classification of epg waveforms of aphid utilizing wavelet kernel extreme learning machine. Applied Artificial Intelligence, 37(1):2214766, 2023. doi: 10.1080/08839514.2023.2214766. URL https://doi.org/10.1080/08839514.2023.2214766.

[10] Wilant A. Van Giessen and D. Michael Jackson. Rapid Analysis of Electronically Monitored Homopteran Feeding Behavior. Annals of the Entomological Society of America, 91(1):145–154, 01 1998. ISSN 0013-8746. doi: 10.1093/aesa/91.1.145. URL https://doi.org/10.1093/aesa/91.1.145.

[11] E. Sarria, Miguel Cid, Elisa Garzo, and Alberto Fereres. Excel workbook for automatic parameter calculation of epg data. Computers and Electronics in Agriculture, 67:35–42, 06 2009. URL 10.1016/j.compag.2009.02.006.

[12] Philippe Giordanengo. Epg-calc: A php-based script to calculate electrical penetration graph (epg) parameters. Arthropod-Plant Interactions, 8, 04 2014. URL 10.1007/s11829-014-9298-z.

[13] Timothy A. Ebert, Elaine A. Backus, Miguel Cid, Alberto Fereres, and Michael E. Rogers. A new sas program for behavioral analysis of electrical penetration graph data. Computers and Electronics in Agriculture, 116:80–87, 2015. ISSN 0168-1699. doi: 10.1016/j.compag.2015.06.011. URL https://www.sciencedirect.com/science/article/pii/S0168169915001684.

[14] Anna Markheiser, Giacomo Santoiemma, Alberto Fereres, Michael Maixner, and Daniele Cornara. Xylfeed – analysing dc-epg waveform variables for european spittlebugs and sharpshooters. 11 2022. URL 10.5073/20221107-091816.

[15] Elisa Garzo, Antonio Jesús Álvarez, Aránzazu Moreno, Gregory P Walker, W Fred Tjallingii, and Alberto Fereres. Novel program for automatic calculation of EPG variables. Journal of Insect Science, 24(3):28, 06 2024. ISSN 1536-2442. doi: 10.1093/jisesa/ieae063. URL https://doi.org/10.1093/jisesa/ieae063.

[16] Quang Dung Dinh, Daniel Kunk, Truong Son Hy, Nalam Vamsi, and Phuong D. Dao. Machine learning for characterizing plant-insect interactions through electrical penetration graphic signal. bioRxiv, 2024. doi: 10.1101/2024.06.10.598170. URL https://www.biorxiv.org/content/early/2024/06/11/2024.06.10.598170.

[17] Frances M. Kimmins and W. Freddy Tjallingii. Ultrastructure of sieve element penetration by aphid stylets during electrical recording. Entomologia Experimentalis et Applicata, 39, 1985. URL 10.1111/j.1570-7458.1985.tb03554.x.

[18] W. Fred Tjallingii and Th. Hogen Esch. Fine structure of aphid stylet routes in plant tissues in correlation with epg signals. Physiological Entomology, 18, 1993. URL 10.1111/j.1365-3032.1993.tb00604.x.

[19] Ernesto Prado and W. Freddy Tjallingii. Aphid activities during sieve element punctures. Entomologia Experimentalis et Applicata, 72, 1994. URL 10.1111/j.1570-7458.1994.tb01813.x.

[20] Jian-Qun Chen, Begonia Martin, Yvan Rahbe, and Alberto Fereres. Early intracellular punctures of two aphids on near-isogenic melon lines with and without the virus aphid transmission (vat) resistance gene. European Journal of Plant Pathology, 103:521–536, 08 1997. doi: 10.1023/A:1008610812437. URL https://doi.org/10.1023/A:1008610812437.

[21] Jaime Jiménez, Elisa Garzo, Javier Alba-Tercedor, Aranzazu Moreno, Alberto Fereres, and G. Walker. The phloem-pd: a distinctive brief sieve element stylet puncture prior to sieve element phase of aphid feeding behavior. Arthropod-Plant Interactions, 14, 02 2020. URL 10.1007/s11829-019-09708-w.

[22] Jaime Jiménez, W. Fred Tjallingii, Aránzazu Moreno, and Alberto Fereres. Newly distinguished cell punctures associated with transmission of the semipersistent phloem-limited beet yellows virus. Journal of Virology, 92 (21):10.1128/jvi.01076–18, 2018. doi: 10.1128/jvi.01076-18. URL https://doi.org/10.1128/jvi.01076-18.

[23] W. Fred Tjallingii. Salivary secretions by aphids interacting with proteins of phloem wound responses. Journal of Experimental Botany, 57(4):739–745, 02 2006. ISSN 0022-0957. doi: 10.1093/jxb/erj088. URL https://doi.org/10.1093/jxb/erj088.

[24] Jonas Van Der Donckt, Jeroen Van der Donckt, Emiel Deprost, and Sofie Van Hoecke. Plotly-resampler: Effective visual analytics for large time series. In 2022 IEEE Visualization and Visual Analytics (VIS), pages 21–25, 2022. doi: 10.1109/VIS54862.2022.00013.

[25] Houtao Deng, George Runger, Eugene Tuv, and Martyanov Vladimir. A time series forest for classification and feature extraction. Information Sciences, 239:142–153, 2013. ISSN 0020-0255. doi: 10.1016/j.ins.2013.02.030.

[26] Mustafa Baydogan, George Runger, and Eugene Tuv. A bag-of-features framework to classify time series. IEEE Transactions on Pattern Analysis and Machine Intelligence, 35:2796–802, 11 2013. doi: 10.1109/TPAMI.2013.72.

