## Appendix for "DiscoEPG: A Python package for characterization of insect electrical penetration graph (EPG) signals"

```
root_dir = 'C:\\Dung\\EPG_project\\data'
dataset = EPGDataset(data_path = root_dir, dataset_name = 'SBA-Rag5', inference = False)
✓ 4.9s

Found 1 datasets: 'SBA-Rag5'.
Loading data ...
SBA-Rag5:   5%|▌         | 2/37 [00:00<00:05,  6.63it/s]
10-6-216-ch1 - Analysis is given up to 28800.00 while the recoring's length is 36000.00.
10-6-216-ch1 - Analysis ends at 28800.0 instead of 36000.0.
10-6-216-ch3 - Analysis is given up to 28800.00 while the recoring's length is 36000.00.
10-6-216-ch3 - Analysis ends at 28800.0 instead of 36000.0.
SBA-Rag5:  11%|█         | 4/37 [00:00<00:04,  7.85it/s]
10-7-16-ch1 - Analysis starts at 0.39 instead of 0.
SBA-Rag5:  22%|██▏       | 8/37 [00:01<00:03,  8.34it/s]
2-8-2016-ch8 - Analysis is given up to 28800.00 while the recoring's length is 36000.00.
2-8-2016-ch8 - Analysis ends at 28800.0 instead of 36000.0.
3-1-2016-ch1 - Analysis starts at 0.12 instead of 0.
SBA-Rag5:  73%|███████▎  | 27/37 [00:03<00:01,  8.95it/s]
6-9-2016-ch7 - Analysis starts at 0.27 instead of 0.
SBA-Rag5:  81%|████████  | 30/37 [00:03<00:00,  8.55it/s]
7-1-2016-ch8 - Analysis is given up to 28800.00 while the recoring's length is 36000.00.
7-1-2016-ch8 - Analysis ends at 28800.0 instead of 36000.0.
SBA-Rag5:  89%|████████▉ | 33/37 [00:03<00:00,  8.93it/s]
7-19-2016-ch3 - No analysis (*.ANA) was found.
SBA-Rag5: 100%|██████████| 37/37 [00:04<00:00,  8.48it/s]
Done! Elapsed: 4.46 s
View guidelines checking log at c:\Dung\EPG_project\ML4Insects/Guideline_check.xlsx
```

Figure 4: DiscoEPG quickly loads the database and describes issues with each recording. Guideline checking are automatically performed.

| | A | B | C | D | E | F | G | H | I | J | K | L |
|---|---|---|---|---|---|---|---|---|---|---|---|---|
| 1 | | The recording is always started with NP | Np is always followed by C or last until the end | E2 is always preceded by E1 | pd waveform is always preceded by C and followed by C or Np | The subphase sequence after pd (pdII-1) should be II-2, II-3 | F is always preceded by C | F is always followed by Np or C | G is always preceded by C | G is always followed by Np or C | E1 is always followed by E1e, E2, C, or Np | The end of each recording must be marked (T, code 99) |
| 2 | **10-6-216-ch1** | True. | True. | True. | True. | True. pd-II-2, pd-II-3 is not presented. | True. F is not presented. | True. F is not presented. | True. G is not presented. | True. G is not presented. | True. | True. |
| 3 | **10-6-216-ch3** | True. | True. | True. | At line 61, pd is not preceeded by C. | True. pd-II-2, pd-II-3 is not presented. | True. F is not presented. | True. F is not presented. | True. | At line 60, G is not followed by C or NP. | True. | True. |
| 4 | **10-7-16-ch1** | Starts with C instead of NP. | True. | At line 134, E2 is not preceeded by E1. | True. | True. pd-II-2, pd-II-3 is not presented. | True. | True. | True. G is not presented. | True. G is not presented. | True. | True. |
| 5 | **10-7-16-ch3** | True. | True. | True. | True. | True. pd-II-2, pd-II-3 is not presented. | True. F is not presented. | True. F is not presented. | True. | True. | True. | True. |

Figure 5: The result of checking 11 guidelines proposed by Garzo et al. [15] presented as a table where each row correspond to a recording.

| | A | B | C | D | E | F | G | H | I | J |
|---|---|---|---|---|---|---|---|---|---|---|
| 1 | **name** | **dataset** | **Interval** | **waveforms** | **s_NP** | **n_NP** | **a_NP** | **m_NP** | **s_C** | **n_C** |
| 2 | **01-10-2018-ch1** | BCOA-Wheat | Hour0-1 | NP, C, pd | 1215,59 | 3 | 405,1967 | 178,44 | 2304,03 | 21 |
| 3 | **01-10-2018-ch1** | BCOA-Wheat | Hour0-2 | NP, C, pd, E1, E2, G | 1677,83 | 5 | 335,566 | 178,44 | 4488,8 | 54 |
| 4 | **01-10-2018-ch1** | BCOA-Wheat | Hour0-3 | NP, C, pd, E1, E2, G | 1677,83 | 5 | 335,566 | 178,44 | 4488,8 | 54 |
| 5 | **01-10-2018-ch1** | BCOA-Wheat | Hour0-4 | NP, C, pd, E1, E2, G | 1677,83 | 5 | 335,566 | 178,44 | 5528,14 | 65 |
| 6 | **01-10-2018-ch1** | BCOA-Wheat | Hour0-5 | NP, C, pd, E1, E2, G | 1821,6 | 6 | 303,6 | 161,105 | 6034,15 | 73 |
| 7 | **01-10-2018-ch1** | BCOA-Wheat | Hour0-6 | NP, C, pd, E1, E2, G | 1821,6 | 6 | 303,6 | 161,105 | 6034,15 | 73 |
| 8 | **01-10-2018-ch1** | BCOA-Wheat | Hour0-7 | NP, C, pd, E1, E2, G | 1821,6 | 6 | 303,6 | 161,105 | 6034,15 | 73 |
| 9 | **01-10-2018-ch1** | BCOA-Wheat | Hour0-8 | NP, C, pd, E1, E2, G | 1821,6 | 6 | 303,6 | 161,105 | 6034,15 | 73 |

(a)

| | A | B | C | D | E | F |
|---|---|---|---|---|---|---|
| 1 | **Variables** | **Description** | **10-6-216-ch1** | **10-6-216-ch3** | **10-7-16-ch1** | **10-7-16-ch3** |
| 2 | **dataset** | Name of the dataset | SBA-Rag5 | SBA-Rag5 | SBA-Rag5 | SBA-Rag5 |
| 3 | **recTime** | Total recording time | 10 | 10 | 8 | 8 |
| 4 | **waveforms** | Observed waveforms | NP, C, pd, E1 | NP, C, pd, G | C, pd, NP, E1, E2, F | NP, C, pd, G |
| 5 | **s_NP** | Sum duration of NP periods | 23624,92969 | 9557,860352 | 8728,269531 | 3704,050049 |
| 6 | **n_NP** | Number of NP periods | 13 | 45 | 23 | 15 |
| 7 | **a_NP** | Average duration of NP periods | 1817,302368 | 212,3968964 | 379,4899902 | 246,9366608 |
| 8 | **m_NP** | Median duration of NP periods | 111,1100006 | 67,69999695 | 63,06999969 | 220,2200012 |
| 9 | **mx_NP** | Max duration of NP | 21974,50977 | 2471,939941 | 6434,759766 | 1086,97998 |
| 10 | **s_Pr** | Sum duration of all probes | 5175,069824 | 19242,14063 | 19972,61914 | 25095,94922 |
| 11 | **n_Pr** | Number of probes | 12 | 44 | 23 | 14 |
| 12 | **a_Pr** | Average probe duration | 431,2558289 | 437,3213501 | 868,3747559 | 1792,567871 |
| 13 | **m_Pr** | Median probe duration | 109,7699966 | 14,82499981 | 32,58000183 | 252,4750061 |

(b)

Figure 6: The parameter calculation result presented in row view (sub-figure a) and column view (sub-figure b).

```
dataset.statistical_analysis('s_NP', [0,1], 'ttest')
```

[50] ✓ 0.2s

```
{'ttest': TtestResult(statistic=0.8875080802511642, pvalue=0.3979022456140333, df=9.0)}
```

Figure 7: An example of performing statistical analysis with DiscoEPG.
